# Dynamic Mechanical Loading Reprograms Meniscus Cell-Derived Extracellular Vesicles to Enhance Their Regenerative Potency

**DOI:** 10.64898/2026.01.10.698807

**Authors:** Catherine Cheung, Zizhao Li, Sung-Han Jo, Se-Hwan Lee, Jina Ko, Su Chin Heo

**Affiliations:** McKay Orthopaedic Research Laboratory, Department of Orthopaedic Surgery University of Pennsylvania 36th Street and Hamilton Walk, Philadelphia, PA, 19104, United States; Department of Bioengineering University of Pennsylvania 210 South 33rd Street, Philadelphia, PA, 19104, United States; Department of Dentistry College of Dentistry, Kyung Hee University, Seoul, Republic of Korea

**Keywords:** Mechanical loading, Extracellular vesicles, Meniscus, Regenerative medicine

## Abstract

The meniscus is a fibrocartilaginous tissue critical for knee stability and load distribution, but its poor healing capacity makes regeneration after injury a major clinical challenge. Extracellular vesicles (EVs) are emerging as promising cell-free therapeutics for tissue regeneration. However, their clinical translation remains limited by low yield and variable bioactivity. Here, we investigated how dynamic mechanical loading regulates the production, composition, and function of meniscus fibrochondrocyte-derived EVs (MFC-EVs). Using a custom bioreactor system, cells were subjected to physiologically relevant cyclic tensile loading. Mechanical stimulation significantly increased EV production and secretion, likely through the ESCRT-independent pathway, without altering vesicle size or tetraspanin expression. Functionally, “mechanically primed” EVs enhanced aggrecan expression in recipient mesenchymal stromal cells (MSCs), mirroring the loading-response phenotype of their source cells. Proteomic profiling identified 380 unique proteins across all EV groups, with loaded EVs enriched in extracellular matrix- and cytoskeleton-associated proteins, as well as pathways related to tissue morphogenesis, cell migration, and cartilage development. Together, these findings demonstrate that physiologic mechanical loading enhances both the yield and regenerative potency of MFC-EVs by enriching their cargo with matrix- and development-associated proteins, providing a scalable and biologically inspired approach for engineering high-efficacy EV therapeutics for meniscus repair and musculoskeletal regeneration.

## INTRODUCTION

The menisci are fibrocartilaginous tissues that provide joint stability, shock absorption, and load distribution in the knee joints [1]. Despite their important role in knee biomechanics, the menisci are highly susceptible to injury due to the complex and repetitive loading environment of the joint. Current nonsurgical treatments such as physical therapy and/or corticosteroid injections primarily aim to alleviate pain rather than promote tissue repair [1,2]. Surgical intervention is typically offered when conservative treatments fail or when patients present with mechanical symptoms such as catching or locking of the knee [1,2]. Although tears localized in the outer one-third of the meniscus, the vascular “red” zone, may heal following repair, the majority of meniscus injuries occur in the avascular inner “white” regions and are treated by arthroscopic partial meniscectomy (APM), in which the damaged portion of tissue is excised. While APM can effectively relieve symptoms, it fails to restore the native biomechanical or biological functions of the meniscus [3,4]. Consequently, approximately two-thirds of patients who undergo meniscus tear treatment develop osteoarthritis (OA) within 5 -15 years after injury, regardless of intervention [5]. Given that OA is a leading cause of disability worldwide, affecting nearly 600 million individuals globally and projected to increase with aging populations [6], developing effective regenerative strategies that prevent OA onset by effectively repairing meniscal injuries remains an urgent clinical need.

The meniscus has a particularly limited healing response largely due to its minimal vascularity and dense extracellular matrix (ECM), which restricts immune cell infiltration, progenitor cell recruitment, and nutrient diffusion [7,8]. Recently, extracellular vesicles (EVs) have emerged as a promising strategy to address these limitations. EVs are nanoscale, membrane-bound vesicles that transport bioactive cargo, including nucleic acids, lipids, proteins, and metabolites, that reflect the physiological state of their cell sources [9,10]. Through paracrine and autocrine signaling, EVs regulate a wide range of biological processes, including cell proliferation [11,12], cell migration [13], differentiation [14], matrix remodeling [15], angiogenesis [16,17], and immune modulation [18,19]. Their small size enables diffusion through dense matrices such as those of cartilage or meniscus. Importantly, compared to direct cell transplantation, EVs have greater stability and lower immunogenicity [9,20,21], positioning them as attractive candidates for next-generation, cell-free regenerative therapeutics. However, the clinical translation of EV-based therapies remains limited by variability in cargo composition and function, which depend on the type and physiological state of their cell source. Moreover, conventional production methods yield low quantities, requiring large-scale cell expansion to achieve therapeutic doses. These challenges highlight the need for strategies that enhance both the yield and bioactivity of EVs for regenerative therapeutics.

In musculoskeletal applications, mesenchymal stromal cell (MSC)-derived EVs have demonstrated therapeutic potential by alleviating inflammation through macrophage repolarization, promoting cartilage matrix synthesis, and delaying OA progression in preclinical models [11,22,23]. Similarly, chondrocyte-derived EVs enhance chondrogenesis in MSC-based engineered cartilage constructs [24]. Despite these advances, little is known about EVs secreted by meniscus fibrochondrocytes (MFCs), the primary resident cells of the meniscus. MFC-derived EVs may mediate intra-meniscal communication and crosstalk with surrounding progenitor cells; however, their production, regulation, and function remain largely unexplored.

The knee joint is a highly dynamic, mechanically active environment that supports weightbearing and mobility. Mechanical stimulation is essential not only for joint development but also for maintaining tissue homeostasis in adulthood [25,26]. Clinically, controlled mechanical loading, such as that achieved through physical therapy, often can yield outcomes comparable to surgical intervention [27–30]. However, the underlying cellular and molecular mechanisms remain poorly defined. Given that EVs are central mediators of intercellular communication [9,10], they may serve as the key effectors coupling mechanical cues to homeostatic and regenerative responses in the meniscus.

Therefore, in this study, we investigated how dynamic mechanical loading influences MFC-EV production and the subsequent effects of these EVs on progenitor cell (e.g., MSC) behavior. To do this, we used a custom bioreactor system to apply cyclic tensile loading to MFC-laden aligned polycaprolactone (PCL) nanofibrous scaffolds, and the secreted EVs were collected from the conditioned media. Mechanical loading significantly increased EV yield without altering their morphology or surface markers. Furthermore, MSCs treated with these “mechanically primed” MFC-EVs displayed gene expression changes, most notably enhanced aggrecan expression, that closely mirrored their EV source, indicating load-dependent signal transmission between cells. Together, these findings identify mechanically primed MFC-EVs as critical mediators of matrix regulation and suggest their promise as a cell-free therapeutic strategy for meniscus repair. This work further indicates the potential of integrating mechanical cues with EV-based approaches to advance biologically inspired therapies for joint regeneration.

## RESULTS AND DISCUSSION

### Dynamic Mechanical Loading Enhances EV Secretion by Meniscus Cells

The meniscus resides in a dynamically active joint, where physiological mechanical loading is essential for maintaining tissue integrity and function. Both overloading and underloading disrupt this delicate balance, resulting in matrix degradation and loss of homeostasis [31,32]. In this context, we sought to determine whether physiologically relevant tensile loading [33] modulates MFC-EV secretion and function. Given that meniscus cells are continuously exposed to mechanical forces during daily activities and that these forces regulate cell behavior and differentiation [34], we first investigated whether dynamic loading influences EV secretion by MFCs and MSCs cultured on aligned PCL nanofibrous scaffolds (**Figure 1a**). Dynamic tensile loading (L) of MFCs increased their EV yield by 5- to 10-fold compared to static free-swelling non-loading (NL) culture conditions across all experiments (**Figure 1b**). To determine whether this effect would be cell type-specific, we applied the same loading regimen to MSCs and observed a 4- to 6-fold amplification of EV secretion relative to NL controls (**Figure 1c**). These results demonstrate that dynamic mechanical stimulation broadly promotes EV secretion in multiple cell types in the knee joint. These findings are consistent with prior reports showing that EV biogenesis and release are mechanosensitive processes influenced by substrate stiffness, shear stress, and mechanical loading [35–37]. As the primary focus of this study was on MFC-EVs, all subsequent analyses were performed using MFC-EVs.

**Figure 1:**
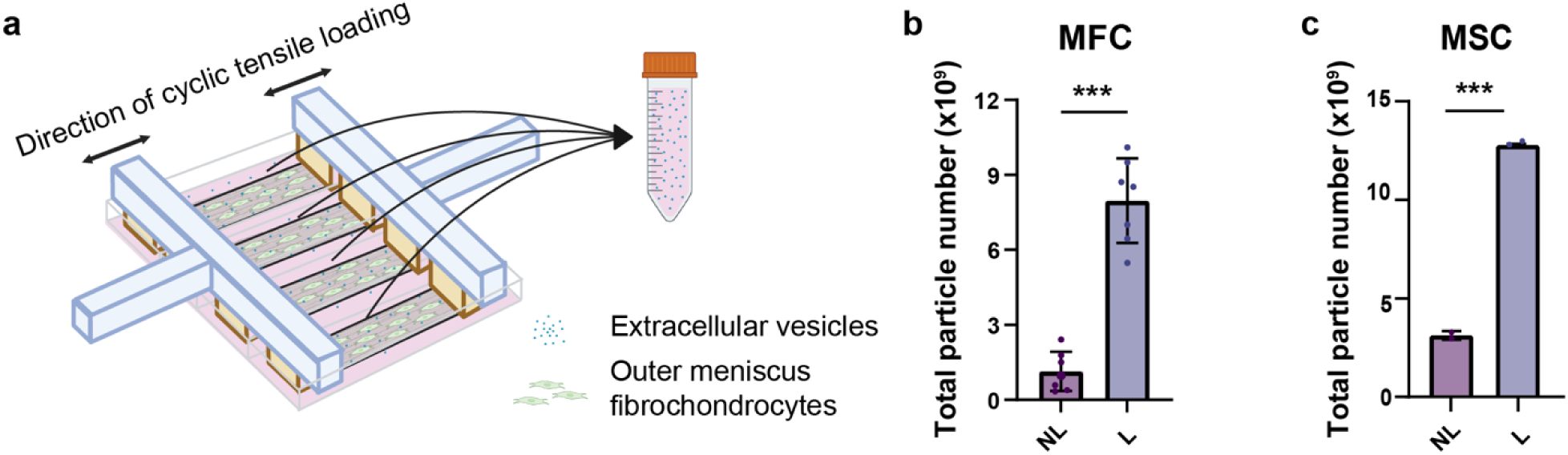
**Dynamic mechanical loading increases EV yield in multiple cell types**. **(a)** Schematic of the custom bioreactor setup and EV collection process; medium is collected from four technical replicate samples per condition and pooled for subsequent EV isolation and analysis. Illustration created with BioRender (https://BioRender.com/5z9hfxz). **(b)** Comparison of EV yield between dynamic loading (L) and non-loading (NL) MFCs (n=7 biological replicates, ***: p<0.001). **(c)** Comparison of EV yield between L and NL MSCs (n=3 biological replicates, ***: p<0.001).

To confirm EV identity and assess morphological characteristics, we next performed multiple EV characterizations. Nanoparticle tracking analysis (NTA) revealed comparable size distributions between static NL control and mechanically primed L EVs, both peaking around 150 nm (**Figure 2a**), consistent with the expected size of exosomes and other small EVs [9,38]. In addition, scanning electron microscopy (SEM) further verified EV morphology, revealing numerous spherical particles smaller than 200 nm (**Figure 2b**). To examine surface marker composition at the single-EV level, we employed stochastic optical reconstruction microscopy (STORM), a super-resolution microscopy technique, with clustering analysis. This approach identified a heterogenous population of EVs containing one, two, or all three of the canonical tetraspanins, CD9, CD63, and CD81 (**Figure 2c-e**). The relative distributions of single-, double-, and triple-positive EVs were not notably different between NL and L groups (**Figure 2d**), indicating that dynamic loading has minimal impact on tetraspanin expression. Similarly, the size distributions of these subpopulations were comparable between NL and L groups (**Figure 2e**), consistent with the NTA results. Taken together, these analyses demonstrate that while dynamic mechanical loading markedly increases EV yield, it does not alter their size, morphology, or surface maker composition.

**Figure 2:**
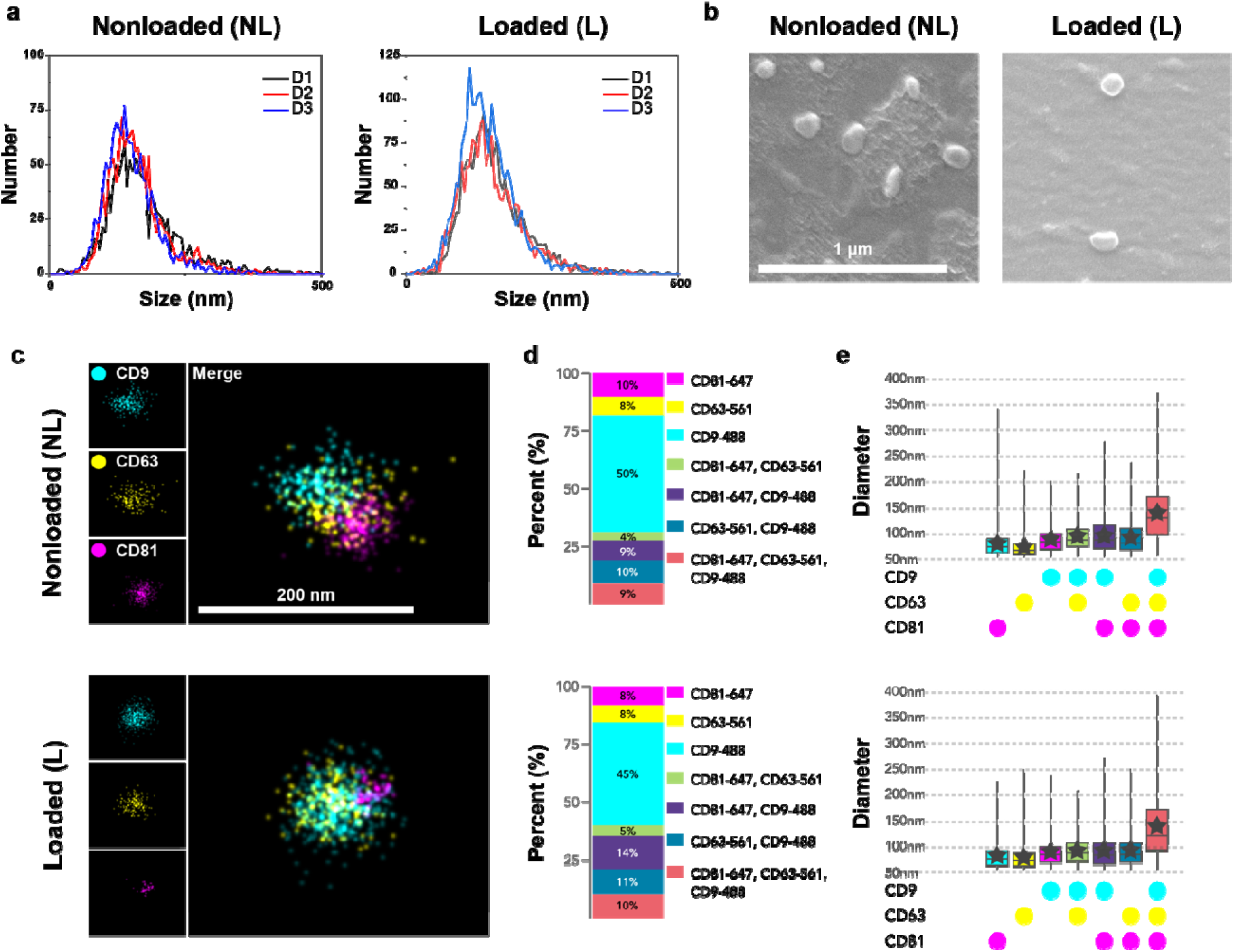
**Characterization of isolated MFC-EVs**. **(a)** Representative size distributions of collected EVs analyzed by nanoparticle tracking analysis (NTA) from 3 different donors (D1-D3). Representative images acquired using **(b)** scanning electron microscopy (SEM) and **(c)** super-resolution microscopy (STORM) of NL-EVs and L-EVs. **(d)** Quantification of STORM clustering analysis showing proportions of EVs positive for individual (CD9, CD63, or CD81), dual, or triple tetraspanin expression. **(e)** Size distributions of collected EVs using STORM imaging and clustering analysis. (NL: n=2,144 EVs, L: n=2,326 EVs).

### Dynamic Loading Increases EV Yield Through ESCRT-Independent Biogenesis

After establishing that dynamic mechanical loading significantly enhances EV secretion, we next sought to elucidate the mechanisms through which this occurs by identifying the pathways responsible for the observed mechano-responsiveness. To this end, several mechanotransduction pathways were pharmacologically inhibited during mechanical loading to identify those contributing to EV production.

Pharmacological inhibition of the Rho/ROCK and actomyosin contractility pathway, which regulates cytoskeletal tension and cell morphology [39], did not affect EV yield under either static free swelling or dynamic loading conditions (**Figures 3a, S1**). Similarly, blocking the YAP/TAZ transcriptional regulators, key mediators of mechanosensitive gene expression [40], did not alter EV secretion (**Figure 3b**). Calcium chelation with BAPTA-AM, used to inhibit intracellular calcium flux as a secondary messenger in mechanotransduction [41], also failed to significantly impact EV output (**Figure 3c**). In contrast, the inhibition of the ESCRT-independent EV biogenesis pathway with GW4869 [42] reduced EV production in non-loading controls, and this reduction could not be fully rescued by dynamic loading (**Figure 3d**). These results indicate that MFCs predominantly utilize ESCRT-independent mechanisms for EV generation, and that dynamic mechanical loading enhances EV yield, at least in part, through this pathway.

**Figure 3:**
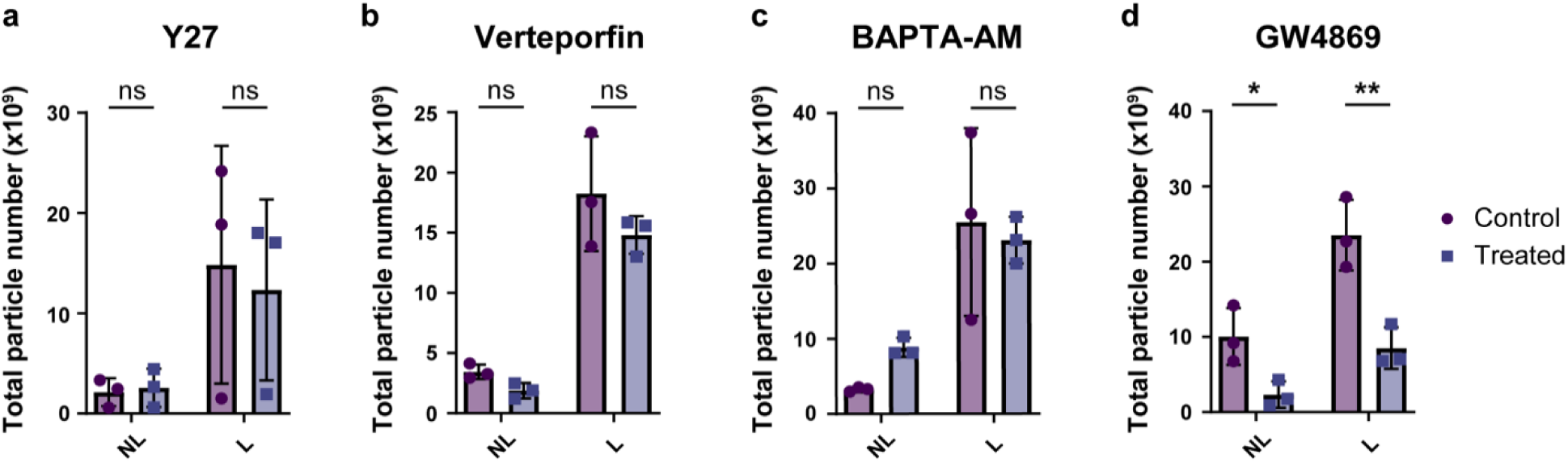
Identification of mechanoresponsive pathways regulating EV secretion in MFCs. Total EV particles quantified by nanoparticle tracking analysis (NTA) under non-loading (NL) and dynamic loading (L) conditions following treatment with pharmacological inhibitors: **(a)** Y27632 (Rho/ROCK inhibitor), **(b)** Verteporfin (YAP/TAZ inhibitor), **(c)** BAPTA-AM (calcium chelator), and **(d)** GW4869 (inhibitor of ESCRT-independent EV biogenesis pathway). All inhibitor studies were conducted using n=3 biological donors, *: p<0.05, **: p<0.01.

Interestingly, our inhibitor experiments indicated that pathways typically associated with mechanotransduction, such as Rho/ROCK, YAP/TAZ, and intracellular calcium signaling pathways, did not significantly affect EV production under loading conditions. This contrasts with a prior study which found that dynamic stretching increased EV secretion by C2C12 muscle progenitor cells partially through YAP/TAZ activity [14]. This discrepancy may arise from cell type-specific mechanosensitivities and different loading regimens, indicating the complexity of EV mechanobiology. Alternative pathways involving other mechanosensitive Rho GTPases (e.g., RAC1, CDC42) [43] or cell membrane deformation driven by cyclic tensile loading may also play roles in mechanically induced EV generation.

### Mechanically Primed EVs Mediate Cell–Cell Crosstalk to Regulate MSC Differentiation

Given that EVs play important roles in cell to cell communication and in regulating recipient cell behavior and differentiation [14], and that mechanical loading enhances EV secretion, we next investigated whether mechanically primed EVs influence MSC differentiation. To this end, EVs were collected from MFCs subjected to dynamic tensile loading for either 1 day (1dL) or 3 days (3dL) and subsequently applied to MSC cultures (**Figure 4a**).

**Figure 4:**
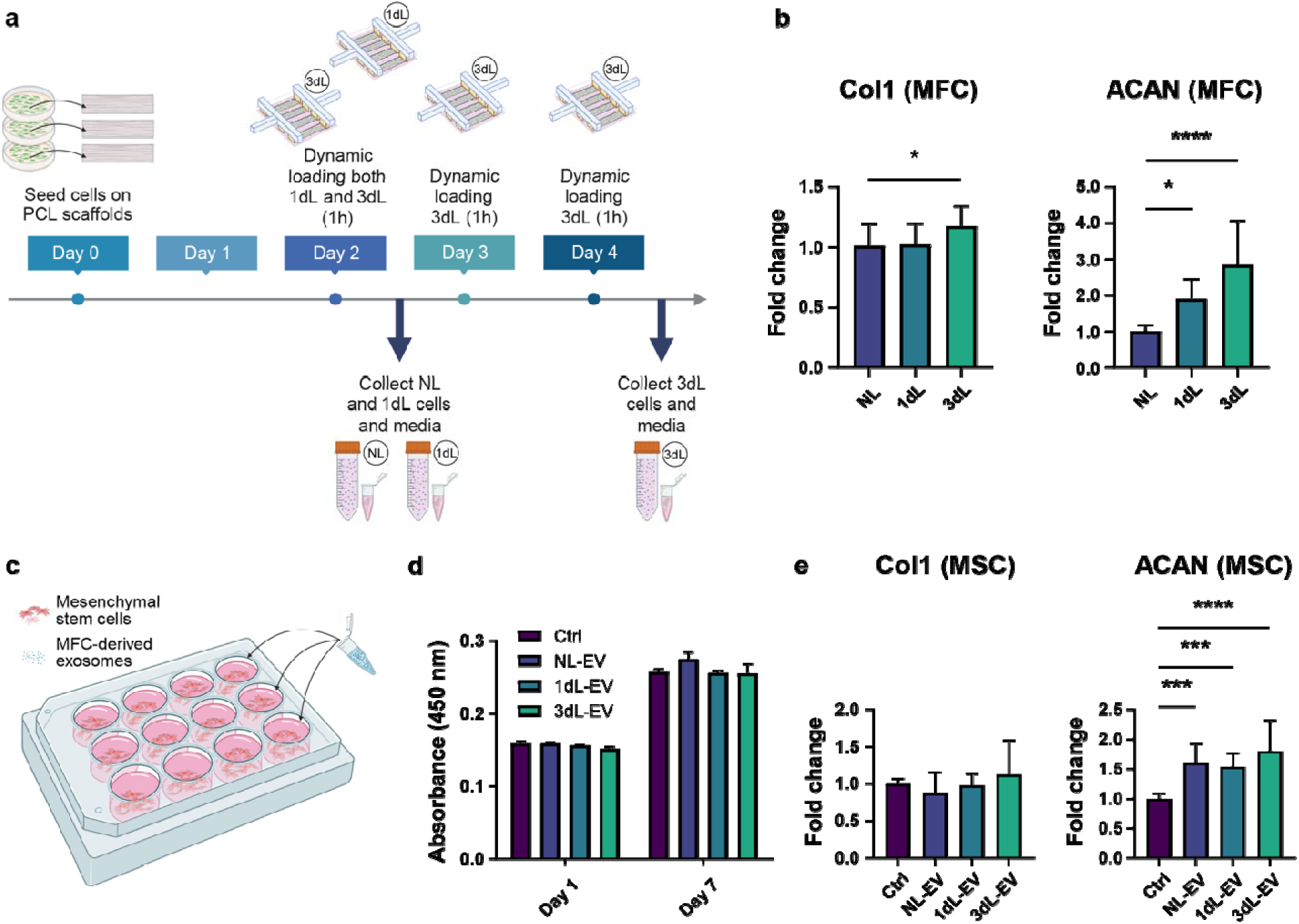
**Mechanically primed MFC-EVs regulate recipient MSC gene expression**. **(a)** Schematic of loading regimen: MFCs were subjected to dynamic cyclic tensile loading for 1 hour either once (1dL) or daily for three consecutive days (3dL). Non-loading (NL) MFCs were maintained under free-swelling conditions for the equivalent duration. Illustration was created with BioRender (https://BioRender.com/es3u9z1). **(b)** Gene expression analysis of NL, 1dL, and 3dL MFCs (n=3 biological replicates, *: p<0.05, ***: p<0.0001). **(c)** Schematic of EV treatment design: MSCs were treated with donor-matched MFC-EVs for 7 days. **(d)** Cell viability and proliferation assays of EV-treated MSCs (n=3 biological replicates). **(e)** Gene expression analysis of MSCs after 7 days of treatment with NL-, 1dL-, or 3dL-EVs (n=3 biological replicates, ***: p<0.001, ****: p<0.0001).

Gene expression analysis of the source MFCs revealed minimal changes in type I collagen (Col1) following 1 hour of daily loading for 1 or 3 days, whereas aggrecan (ACAN) expression increased significantly with prolonged loading (**Figure 4b**). To assess whether EVs secreted under these conditions could impact recipient progenitor cells, MSCs were seeded and treated with the EVs collected from non-loading, 1dL, and 3dL conditions (NL-EVs, 1dL-EVs, and 3dL-EVs, respectively) (**Figure 4c**). CCK-8 assays showed no differences in MSC viability or proliferation after seven days of treatment (**Figure 4d**), confirming the cytocompatibility of all three EV types. Notably, MSCs treated with mechanically primed EVs exhibited gene expression profiles that paralleled those of their source MFCs, showing little change in type I collagen (Col1) but significantly higher aggrecan (ACAN) expression compared to control (**Figure 4e**).

These data demonstrated that mechanical loading not only increased EV yield but also produced mechanically primed EVs capable of regulating gene expression in recipient MSCs. Specifically, these EVs induced a significant upregulation of aggrecan, a key glycosaminoglycan responsible for the compressive strength of fibrocartilaginous tissues, while maintaining similar trends in type I collagen expression. This pattern closely mirrored that of the source MFCs under applied loading, suggesting that EVs may help transmit load-dependent biochemical signals from MFCs to surrounding progenitor cells. Together, these results indicate that EVs secreted by mechanically primed MFCs promote differentiation of MSCs toward a matrix-producing, fibrocartilaginous phenotype reflective of the source cells.

### Dynamic Loading Reprograms EV Protein Cargo Toward Regenerative and Matrix-Associated Functions

Given that dynamic mechanical loading influences EV secretion by MFCs, which regulate differentiation in recipient MSCs, we next examined whether mechanical loading altered the protein composition of the mechanically primed EVs. To do this, EV proteins were isolated and analyzed by mass spectrometry following SDS-PAGE separation and stabilization (**Figure 5a**). Compared with cell-free media controls (NL-neg and 3dL-neg), both NL-EVs and 3dL-EVs displayed distinct and more abundant protein bands (**Figure 5a**). After subtracting media-derived contaminant proteins, a total of 380 unique proteins were identified in both NL-EVs and 3dL-EVs (**Figure 5b**). Among these, 48 proteins (12.6% of total) were unique to NL-EVs, 153 proteins (40.3% of total) were unique to 3dL-EVs, and 179 proteins (47.1% of total) were shared between the two conditions (**Figure 5b**). The top 30 proteins in each group accounted for more than 80% of their total protein content (**Figure 5c**). Representative top 20 proteins in NL-EVs, 3dL-EVs, and those upregulated in 3dL-EVs relative to NL-EVs (Upreg.) are shown in **Figure 5d**, with a comprehensive list provided in **Supplementary Table 1**. We found that dynamic mechanical loading increased the abundance of proteins such as COL14A1, DCN, ANXA1, ACTB, HRG, COL1A1, COL1A2, TNC, NELL2, and DYNC1H1 within EVs (Upreg. In **Figure 5d**), highlighting the enrichment of extracellular matrix-, cytoskeletal-, and regeneration-associated factors that may promote meniscal repair. Protein-protein interaction analysis demonstrated strong interconnections among the top proteins in NL-EVs, 3dL-EVs, and the most upregulated proteins (**Figure 5e**). Gene Ontology (GO) enrichment of the top 30 proteins in each group showed that both NL-EVs and 3dL-EVs were enriched for biological processes and molecular functions related to negative regulation of catalytic and endopeptidase activity, as well as collagen-associated cellular components (**Supplementary Figures S3, S4**). Notably, proteins upregulated in 3dL-EVs were enriched for biological processes associated with cell migration, tissue development, and cartilage development (**Figure 5f**), and for molecular functions involving glycosaminoglycan (GAG) binding (**Figure 5h**). Cellular components of these proteins were primarily linked to collagen organization and the extracellular matrix (**Supplementary Figure S2**).

**Figure 5:**
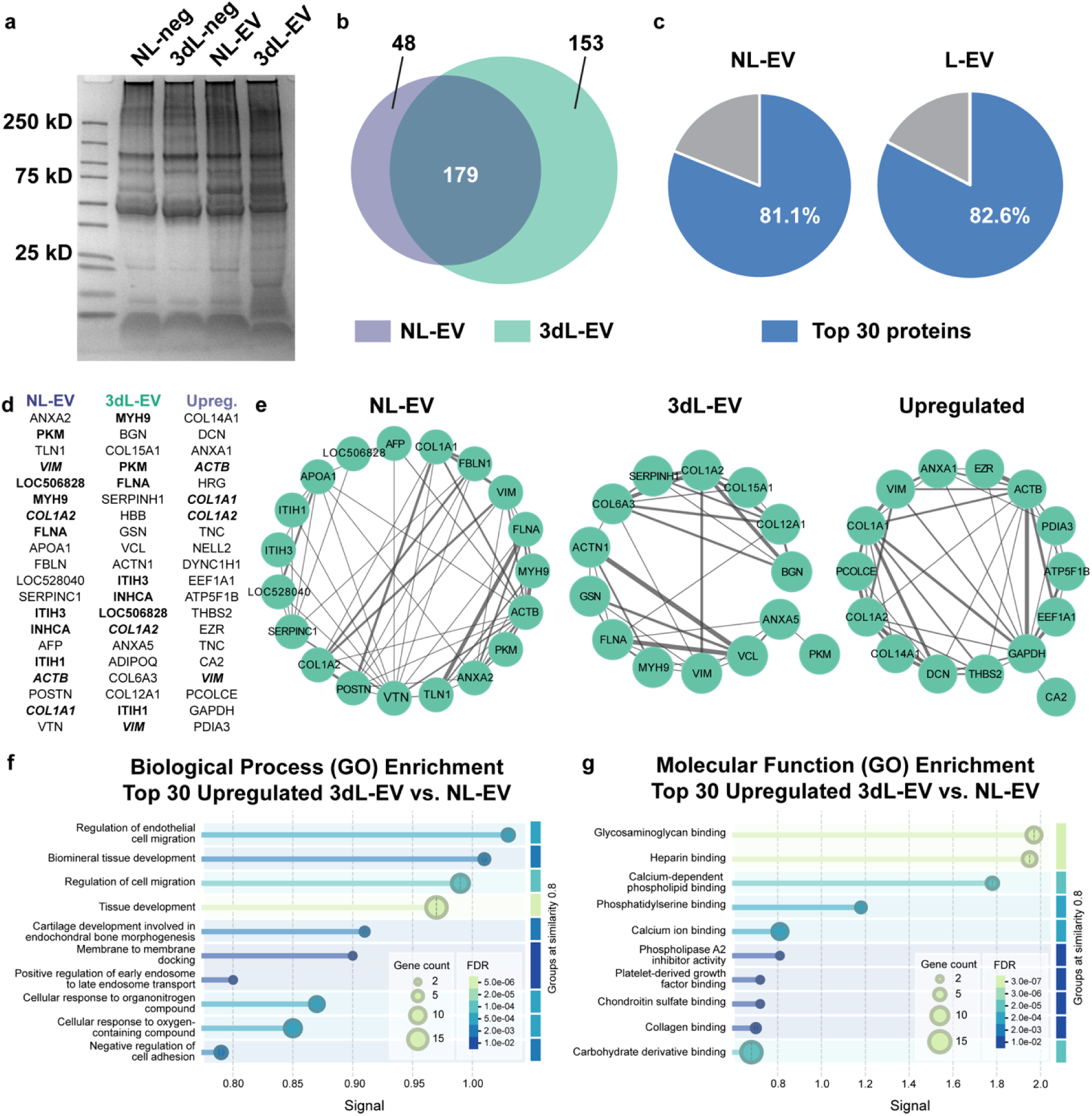
**Proteomic profiling of MFC-derived EVs using liquid chromatography-tandem mass spectrometry (LC-MS/MS)**. **(a)** SDS–PAGE analysis of samples prepared for proteomics compared with a Bio-Rad protein ladder. Groups (left → right): non-loading without cells (NL-neg), loading without cells (3dL-neg), non-loading with cells (NL-EV), and loading with cells (3dL-EV). (b) Venn diagram showing overlapping and unique proteins identified in NL- and 3dL-EVs. (c) Relative proportion of the top 30 proteins in NL-EVs (left) and 3dL-EVs (right). (d) Top 20 proteins detected in NL-EVs, 3dL-EVs, and those upregulated in 3dL-EVs compared with NL-EVs. Bold indicates proteins shared between NL- and 3dL-EVs; bold-italics indicate proteins also found among the top upregulated proteins. (e) Protein–protein interaction (PPI) networks of the top proteins in NL-EVs, 3dL-EVs, and the most upregulated proteins in 3dL-EVs. (f) Gene Ontology (GO) enrichment of the top 30 upregulated proteins highlighting enriched biological processes, and (h) molecular functions, where dot size and color represent gene counts and false-discovery rates (FDRs), respectively.

Together, our proteomic analysis revealed that mechanical priming profoundly altered EV composition. 3dL-EVs contained broader and more diverse proteins than NL-EVs, indicating that mechanical loading enhances both the quantity and regenerative quality of MFC-EVs by enriching cargo proteins linked to matrix remodeling and fibrocartilage formation. Proteins unique to 3dL-EVs are collectively linked to collagen fibrillogenesis, ECM stabilization, cytoskeletal remodeling, and anti-inflammatory signaling, all of which are key processes for tissue repair. Moreover, the GO analysis confirmed that upregulated proteins were enriched for biological processes such as cell migration, tissue morphogenesis, and cartilage development, as well as molecular functions related to glycosaminoglycan binding and ECM interactions. These data support that mechanical loading enhances both the quantitative yield and qualitative potency of EVs as bioactive mediators of matrix remodeling and fibrocartilage regeneration, underscoring their potential as bioactive mediators for enhanced meniscus repair and tissue homeostasis.

From a translational perspective, our study presents a feasible strategy to overcome one of the primary barriers in EV-based therapeutics: their inherently low yield. Dynamic mechanical loading offers a simple and physiologically relevant approach to boost EV production without requiring genetic or chemical manipulation. These findings corroborate other studies that show that modulation of the physical microenvironment can optimize EV output and cargo composition [37]. Importantly, we showed that this strategy also may augment the therapeutic potential of EVs, especially when they are derived from mechanosensitive cell types such as meniscus cells, chondrocytes, or tendon fibroblasts. Our approach may be scalable for generating high-potency EV preparations for musculoskeletal repair and regeneration.

Despite these advances, there are certain limitations of this study. Firstly, MSC treatment assays were conducted in 2D culture, which is poorly representative of their native environment. Future studies should evaluate EV-mediated effects in 3D scaffold or organoid systems that better mimic *in vivo* conditions. Additionally, examining different loading conditions including amplitudes, frequencies, and durations could reveal optimal parameters for maximizing EV yield and regenerative efficacy. Finally, *in vivo* validation in meniscus injury models will be necessary to confirm the therapeutic potential of mechanically primed EVs and to assess their safety, retention, and bioactivity in joint environments.

## CONCLUSION

EVs have emerged as promising tools for regenerative therapies due to their ability to mediate intercellular communication, deliver bioactive cargo, and promote tissue repair. However, their clinical translation remains limited primarily by low production yield and variability in bioactivity. This study establishes that physiological mechanical loading of MFCs regulates both the biogenesis and functional content of MFC-EVs. We demonstrated that dynamic mechanical loading significantly improves MFC-EV quantity through ESCRT-independent mechanisms and simultaneously enriches their cargo with proteins associated with ECM organization and tissue development. These “mechanically primed” EVs transmit these load-dependent cues to promote recipient MSC differentiation, highlighting their potential as bioactive, cell-free therapeutics for meniscus repair. Together, our findings provide new mechanistic insight into EV-mediated mechanobiology and present a scalable, biologically inspired approach for engineering next-generation regenerative therapies.

## EXPERIMENTAL SECTION

### Primary cell isolation

Fresh juvenile (∼3 months) bovine stifle joints (Research 87) were cleaned and the menisci detached from the tibial plateau with a scalpel. MFC isolation was carried out as previously described but only from the outer zone [8,44]. Briefly, the outer one-third of the tissue was cut from the remaining tissue, chopped into small cubes (2-4 mm), placed sparsely into tissue culture dishes, and covered in basal medium (BM) consisting of Dulbecco’s Modified Eagle’s Medium (DMEM, Gibco) supplemented with 10% fetal bovine serum (FBS, Gibco) and 1% penicillin-streptomycin (Corning). Media were changed every 2-3 days. Cells began migrating out of tissue fragments after 7-10 days of culture, and the remaining tissue fragments were removed after ∼14 days. For subsequent use, P0 cells were expanded in BM and either frozen or passaged to P1.

MSCs were isolated from the bovine stifle joint as previously described [45,46]. Briefly, the femur and tibia were sectioned using a reciprocating saw to expose the trabecular bone. Trabecular bone fragments were removed and submerged in dissection media comprised of DMEM supplemented with 2% penicillin-streptomycin and 1 g heparin/500 mL media. The bone-medium mixture was shaken, and the supernatant was collected into a clean conical tube. The tubes of media were centrifuged, and the resulting pelleted cells were resuspended in BM and plated in tissue culture dishes. Media was changed every 2-3 days. Cells were initially frozen at P0 once they reached 90% confluency. For subsequent use, P0 cells were expanded in BM and either frozen or passaged to P1.

### Preparation of nanofibrous scaffolds

Electrospun nanofibrous polycaprolactone (PCL) scaffolds were prepared as described previously [47–49]. Briefly, PCL was dissolved at 35% w/v in a 1:1 (v/v) mixture of N,N-dimethylformamide (DMF) and tetrahydrofuran (THF). A total of 11 mL of PCL solution was loaded into a 20 mL syringe fitted with an 18-guage blunt-tip needle and electrospun onto a rotating and grounded mandrel at a flow rate of 2.5 mL/hour. A high voltage of ∼17 kV was applied to the solution during electrospinning. To maintain alignment of the collected nanofibers, the surface speed of the mandrel was set at ∼10 m/s. The resulting PCL sheet was cut into 21 mm x 55 mm strips with the long side parallel to the prevailing fiber direction. Prior to use PCL scaffolds were rehydrated in decreasing concentrations of ethanol (100%, 70%, 50%, and 30%) for 30 minutes each, washed in PBS for 30 minutes, and coated in fibronectin (Sigma) overnight at room temperature. Scaffolds were sterilized for 30 minutes under UV light prior to cell seeding.

### Bioreactor loading

Passage 2 MFCs were expanded in tissue culture dishes, trypsinized, and seeded onto scaffolds at a density of ∼8 × 10^6^ cells/scaffold in BM. After 2 days of static free swelling culture, scaffolds were transferred to fresh culture dishes containing DMEM supplemented with 10% exosome-depleted FBS (Gibco) and 1% penicillin-streptomycin (Exo-D BM). Loaded scaffolds were affixed into the custom-made bioreactor [46,50,51] and subjected to cyclic tensile loading at 3% strain amplitude and 1 Hz frequency. For characterization and inhibitor studies, the scaffolds underwent 3 consecutive hours of loading. For 1-day and 3-day loading experiments, each round of loading consisted of 1 hour of loading followed by 3 hours of free swelling. In the 3-day loading groups, scaffolds were removed from the bioreactor and returned to the same culture medium overnight between rounds, for a total of three loading cycles (**Figure 4a**). Static (NL) controls remained in free-swelling conditions for the duration of the loading period. At the completion of loading, culture medium was collected for further processing. For RNA extraction, the central region of each scaffold was cut with a scalpel into small (∼2-3 mm) squares and placed into TRIzol reagent for subsequent processing.

### EV isolation and characterization

Culture medium collected from scaffolds was first centrifuged at 500 x g for 5 minutes at 4°C. The supernatant was transferred to a clean tube and centrifuged again at 2000 x g for 20 minutes at 4°C. The resulting supernatant was transferred to 38.5 mL ultra-clear open-top tubes (Beckman) and ultracentrifuged twice at 100,000 × g (24,200 rpm) for 70 minutes at 4 °C using an SW Ti 32 rotor in a Beckman Coulter Optima XPN-80 Ultracentrifuge. After each ultracentrifugation step, the supernatant was carefully removed using serological and micropipettes. The final EV pellet was resuspended in ∼200-500 µL of PBS.

Isolated EVs were characterized by nanoparticle tracking analysis (NTA) using the ZetaView PMX220 Twin. Briefly, resuspended EVs were further diluted in 2 mL of PBS to achieve a density of particles on the order of 10^5^-10^10^ particles per milliliter. 1 mL of the diluted sample was injected into the cell assembly at 25 °C. The instrument then recorded a video from which the particles’ Brownian motion was analyzed to determine their size and concentration. Total EV quantities were calculated using these particle concentrations and the volume of the initial resuspension after ultracentrifugation.

EV morphology was confirmed by scanning electron microscopy (SEM). Briefly, EV suspensions were mixed with an equal volume of 8% glutaraldehyde and incubated overnight at 4 °C to fix. The EVs were then transferred into a 100 kDa centrifugal filter tube, washed with sterile PBS, and centrifuged at 2,000 x g for 3 minutes. Following this, the EVs were dehydrated with increasing concentrations of ethanol (80%, 90%, and 100%). For each ethanol concentration, approximately 10-15 mL of ethanol was added to the filter tube and centrifuged at 2,000 x g for 3 minutes and the filtrate was discarded after each step. The dehydrated samples were resuspended to a final volume of 1 mL and transferred to a clean cover glass to dry. The EVs were sputter coated with iridium using an EMS/Quorum Q150T ES sputter coater and imaged on the FEI Quanta FEG 250 scanning electron microscope.

Stochastic Optical Reconstruction Microscopy (STORM) was performed using the Oxford Nanoimager to visualize EV size and distribution of surface markers CD9, CD63, and CD81 (ONI). Samples were prepared according to the EV Profiler Kit 2 manufacturer’s instructions and co-stained with the antibodies prior to imaging. Images were acquired and analyzed using the AutoEV and CODI platforms.

### Inhibitor treatments

To assess the effects of pathway-specific inhibitors on EV production, scaffolds were prepared and seeded as above. To inhibit the Rho/ROCK signaling pathway, Y27632 (10 µM) was added to scaffolds in Exo-D BM for 1 hour prior to loading. To inhibit actomyosin contractility, blebbistatin (50 µM) was added to scaffolds in Exo-D BM for 1 hour prior to loading. To inhibit YAP/TAZ signaling, verteporfin (1 µM) was added to scaffolds in Exo-D BM immediately prior to loading. To inhibit calcium signaling, calcium chelator BAPTA-AM (25 µM) was added to scaffolds in Exo-D BM for 30 minutes prior to loading. To inhibit ESCRT-independent EV biogenesis, GW4869 (20 µM) was added to scaffolds in Exo-D BM 24 hours prior to loading. All inhibitor studies were conducted using the same loading conditions of 3 hours of cyclic tensile loading at 3% strain amplitude and 1 Hz frequency. All respective static controls were maintained in identical media conditions containing the respective inhibitors for the same durations as the loaded groups. All inhibitors were purchased from Sigma.

### EV treatment of MSCs

MSCs were expanded in 15 cm tissue culture dishes, then seeded into 12-well tissue culture plates at a density of 2 × 10^5^ cells/well in BM. 1 day after seeding, the media was replaced with a chemically defined medium (CM-) consisting of DMEM, dexamethasone (100 µM), ascorbate-2-phosphate (50 µg/mL), L-proline (40 µg/mL), sodium pyruvate (100 µg/mL), ITS+ premix (1X, Corning), penicillin-streptomycin (1%), bovine serum albumin (1.25 mg/mL), and linoleic acid (5.3 µg/mL). For EV treatments, 1 × 10^4^ EVs/cell (based on the cell seeding density) was added to each well (**Figure 4c**). Cell viability was assessed using the CCK-8 reagent according to the manufacturer’s protocol. For gene expression analysis, EV treatments were carried out over 7 days, with an EV-containing media change on day 3. On 7 days, the cells were lysed in TRIzol reagent for RNA extraction.

### Gene expression analysis with real-time quantitative PCR

Scaffolds or cells in TRIzol underwent RNA extraction using Zymo RNA mini-prep columns (Zymo Research, Irvine, CA), according to the manufacturer’s protocols. Extracted RNA was converted to cDNA using the ProtoScript II Reverse Transcriptase kit (NEB) using random primers according to manufacturer’s protocols. Real-time quantitative PCR (RT-qPCR) was carried out using SYBR Green Master Mix (Applied Biosystems) according to manufacturer’s protocols. Primer sequences for RT-qPCR are listed in Table 1. GAPDH was used as the housekeeping reference gene, and relative gene expression levels were calculated using the ΔΔCT method.

**Table 1:**
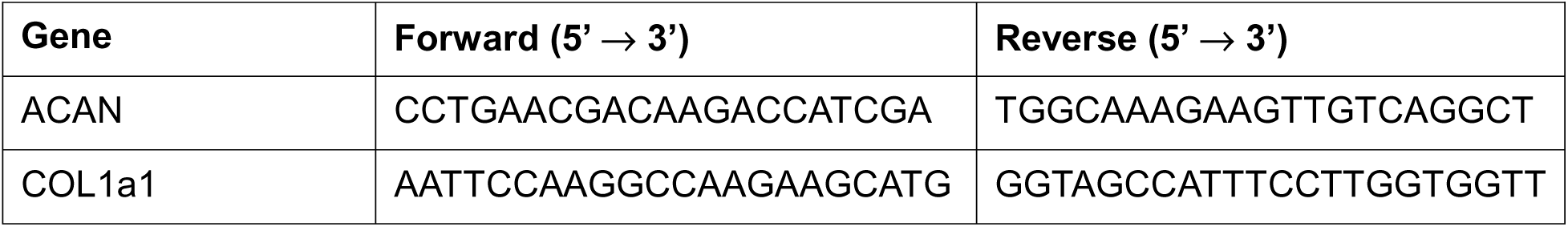

### Proteomic analyses

For proteomic analysis, EVs were isolated using polyethylene glycol (PEG) precipitation as previously described [52]. To prepare PEG solution, 10 kDa PEG was reconstituted in deionized water at a 50% w/v ratio until a homogenous solution was achieved. The solution was sterile-filtered through a 0.22 um filter and used as the stock solution. Cell culture media was collected from both cell-free (negative controls) and cell-containing scaffolds, which were seeded and loaded either once or over three days as described. The media was centrifuged at 500 x g for 5 minutes followed by 2000 x g for 20 minutes to clear cell debris. Then, 8 mL of each media sample was mixed with 2 mL of PEG stock solution and incubated at 4 °C overnight. The following day, the samples were centrifuged at 3,000 x g for 10 minutes at 4 °C. The supernatant was carefully decanted and aspirated from the pellet, which was then resuspended in 50 µL RIPA buffer and vortexed to lyse proteins. Protein concentrations were analyzed using bicinchoninic acid (BCA) protein assay (Thermo) according to manufacturer’s protocols. Equal amounts of protein (18 µg) per sample were prepared in 4x Laemmli reducing sample buffer, boiled for 5 minutes at 105 °C for protein denaturation, and run on a precast 4-20% polyacrylamide gel (Bio-Rad). For SDS-PAGE analysis, samples were run in adjacent wells at 100 V for approximately 1 hour. Gels were then rinsed with deionized water, stained with Coomassie blue for 1 hour at room temperature, washed with deionized water, and destained overnight. Protein bands were visualized using a Gel Doc imaging system.

For mass spectrometry, samples were run with a minimum of 3 wells of separation to minimize crossover contamination at 100 V for approximately 30 minutes. Protein bands were then fixed in a solution of methanol and acetic acid. The fixed gels were stained with Coomassie blue at 4 °C for 1 day and subsequently destained for 1 day. After destaining, the protein bands were carefully cut from the gel and placed into a 2 mL microcentrifuge tube filled with deionized water. Samples were then shipped to the Taplin Mass Spectrometry Facility at Harvard University for liquid chromatography-tandem mass spectrometry (LC-MS/MS). Media proteins found in the static and mechanically primed negative controls were subtracted from the cell-containing samples by their intensity percentages. Protein networks of the top 20 expressed and upregulated proteins were visualized and annotated using the Cytoscape tool (version 3.10.4, Cytoscape Consortium 2025) [53]. Gene ontology (GO) enrichment analyses were conducted using the STRING web tool (version 12.0, http://string-db.org, String Consortium 2025) to identify biological process, molecular function, and cellular component terms with a false discovery rate (FDR) <0.05.

### Statistical analysis

All numerical data are reported as the mean and standard deviation of included samples. Three to four biological replicates were used in all studies. Two-sample t tests and one-way analysis of variance (ANOVA) was performed on data, with Tukey’s *post hoc* testing as applicable. Statistical significance was reached at p<0.05. All statistical analyses were carried out on Graphpad Prism 10.

## Acknowledgements

This research was supported by grants from the National Institutes of Health (R01 AR079224, R01 HL163168, P50 AR080581) and the NSF Science and Technology Center for Engineering Mechanobiology (CMMI-1578571).

## Data Availability Statement

The data that support the findings of this study are available from the corresponding author upon reasonable request.

## Supplementary Information

**Figure S1:**
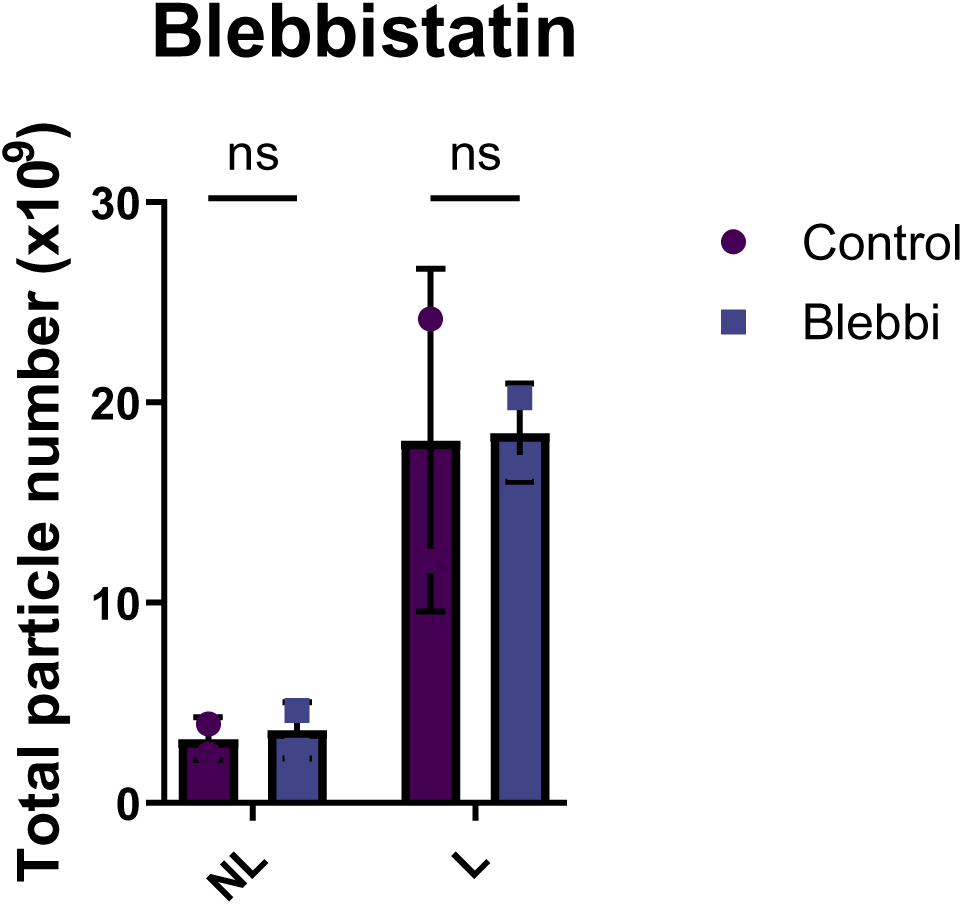
Total EV yield from non-loading (NL) and dynamic loading (L) MFCs treated with blebbistatin(−), an inhibitor of actomyosin contractility (n=2 biological replicates).

**Figure S2:**
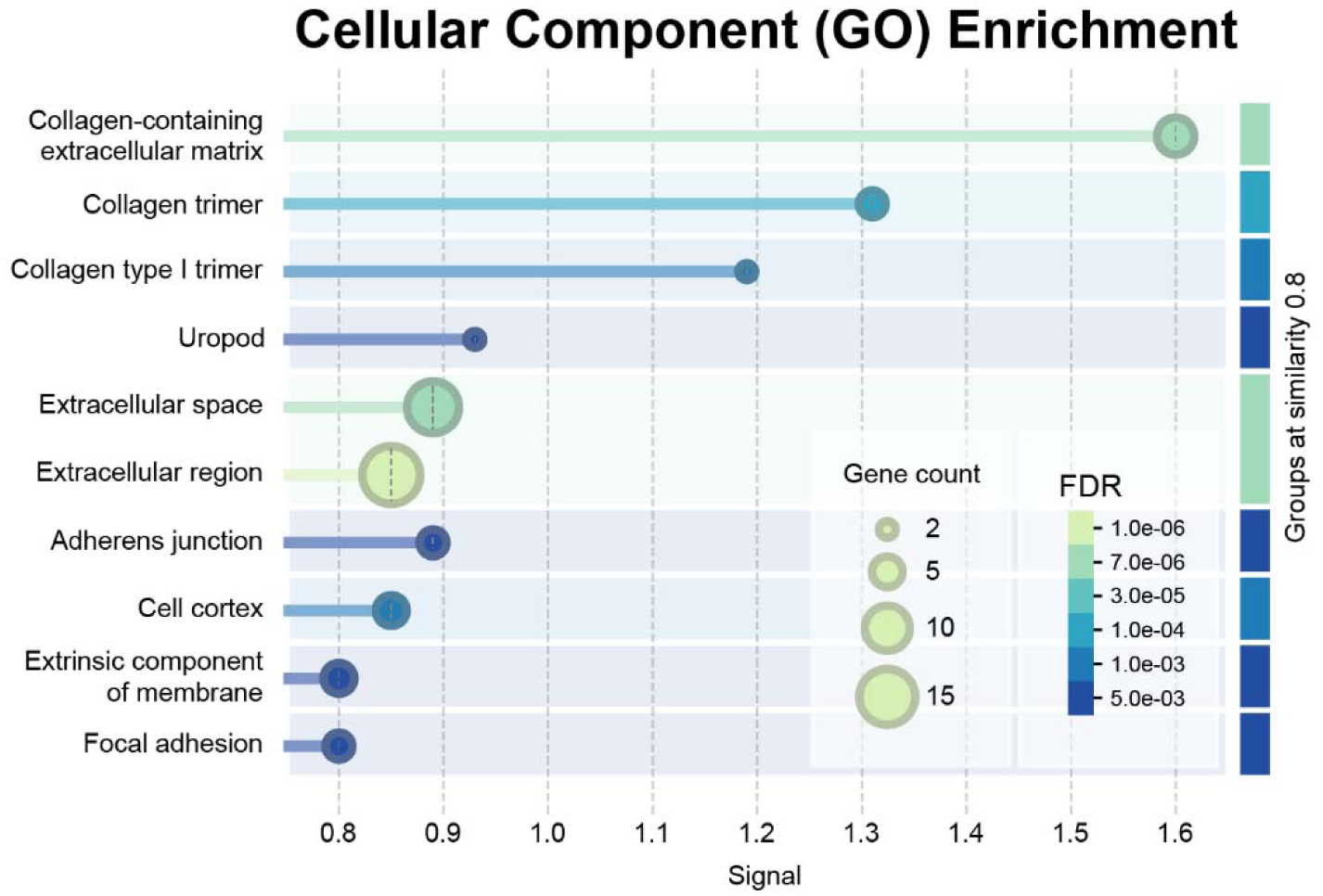
Gene Ontology (GO) enrichment analysis of the top 30 upregulated proteins in dynamic mechanical loading (3dL) EVs compared to non-loading (NL) EVs.

**Figure S3:**
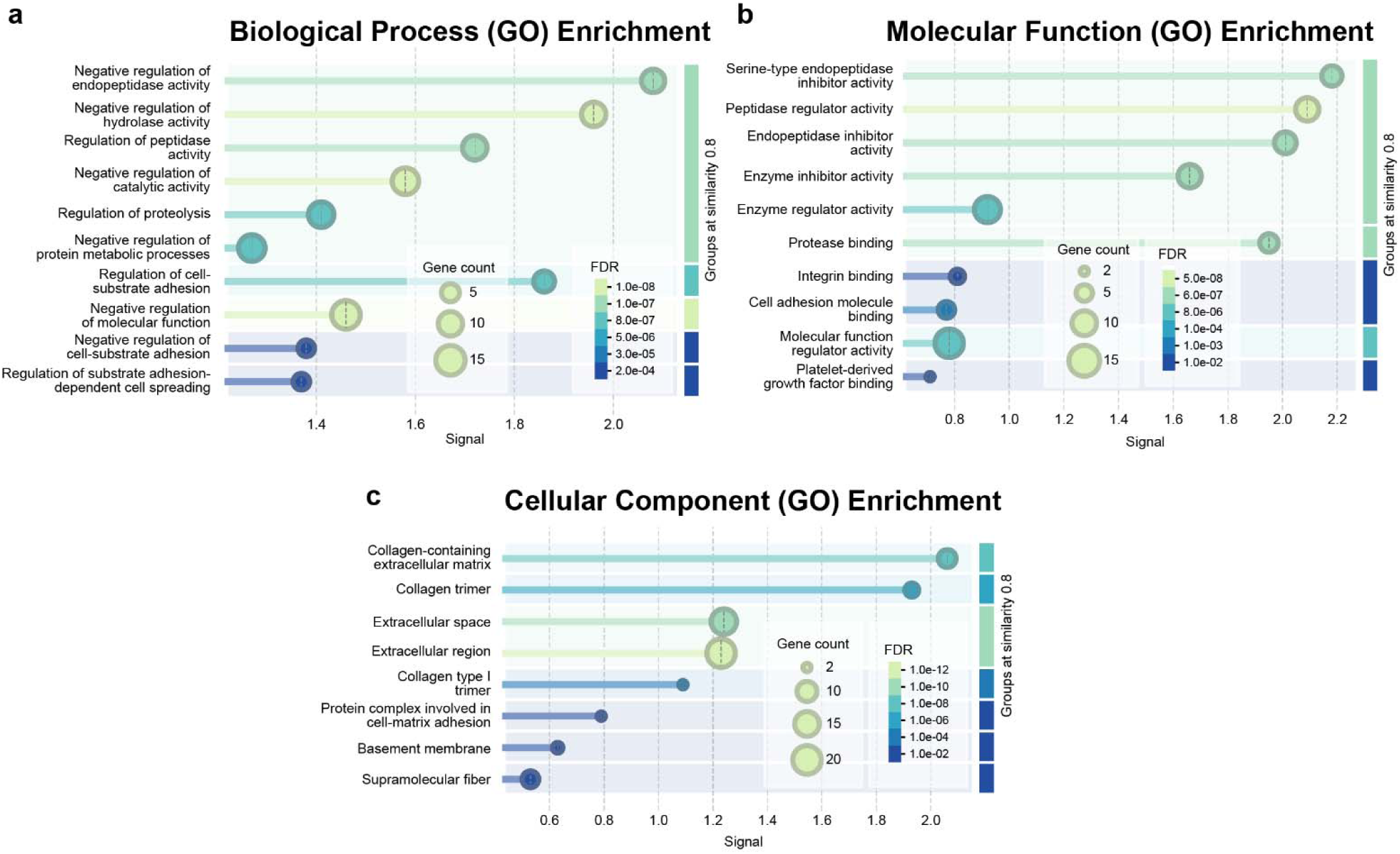
Gene Ontology (GO) enrichment analyses of the top 30 proteins identified in non-loading (NL) EVs. Analyses include (a) biological processes, (b) molecular functions, and (c) cellular components.

**Figure S4:**
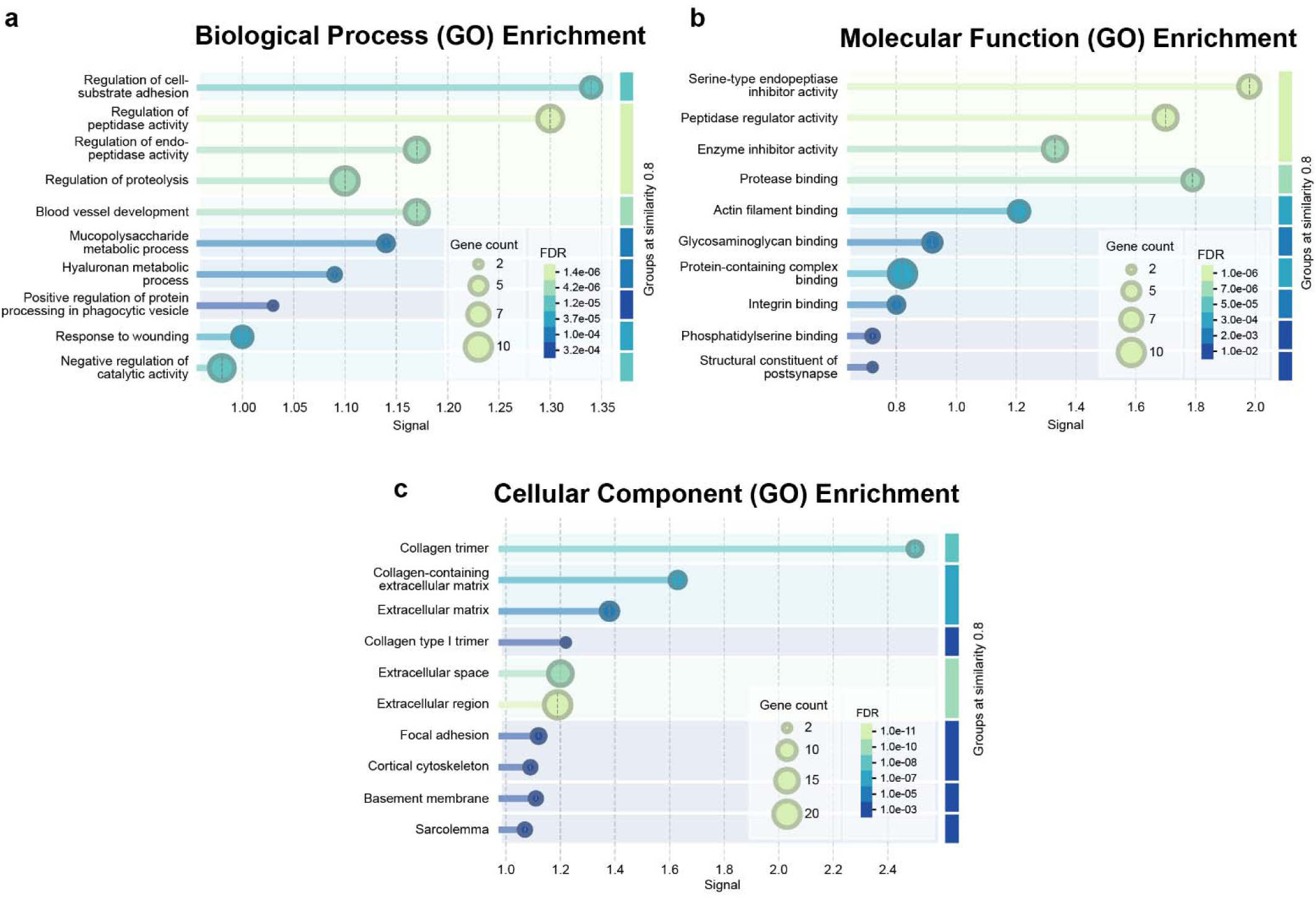
Gene Ontology (GO) enrichment analyses of the top 30 proteins identified in mechanically loaded (3dL) EVs. Analyses include (a) biological processes, (b) molecular functions, and (c) cellular components.

**Supplementary Table 1:** Representative top 50 proteins identified by liquid chromatography–tandem mass spectrometry (LC–MS/MS).

| Top 50 proteins in NL-EVs | Top 50 Proteins in 3dL-EVs | Top 50 proteins upregulated in 3dL-EVs vs. NL-EVs | Top 50 proteins only found in 3dL-EVs |
| --- | --- | --- | --- |
| PLG | TNC | CDH1 | FN1 |
| A2M | PLG | MFGE8 | GSN |
| TNC | MFGE8 | ANXA6 | EDIL3 |
| ADIPOQ | FN1 | LMNA | ALDOA |
| ITIH2 | COL1A1 | ANXA5 | ITIH4 |
| LOC506828 | ACTB | ANXA4 | MMP2 |
| ACTN4 | ACTN4 | ANXA2 | NID1 |
| HBB | ANXA2 | GPI | SERPINE1 |
| COL6A3 | ITIH2 | MSN | VCAN |
| COL12A1 | A2M | SERPINH1 | HTRA1 |
| VTN | VIM | PDIA3 | MST1 |
| COL1A1 | ITIH1 | GAPDH | NT5E |
| POSTN | COL12A1 | PCOLCE | ALDOC |
| ACTB | COL6A3 | VIM | MYO1C |
| ITIH1 | ADIPOQ | CA2 | ATP1A1 |
| AFP | ANXA5 | TNC | ANXA11 |
| INHCA | COL1A2 | EZR | PGK1 |
| ITIH3 | LOC506828 | THBS2 | ACAN |
| SERPINC1 | INHCA | ATP5F1B | CLEC3B |
| LOC528040 | ITIH3 | EEF1A1 | PNP |
| FBLN1 | ACTN1 | DYNC1H1 | FSCN1 |
| APOA1 | VCL | NELL2 | ITGAV |
| FLNA | GSN | TNC | GNAI2 |
| COL1A2 | HBB | COL1A2 | RTN4 |
| MYH9 | SERPINH1 | COL1A1 | YWHAH |
| LOC506828 | FLNA | HRG | LOC784932 |
| VIM | PKM | ACTB | SLC1A5 |
| TLN1 | COL15A1 | ANXA1 | ITGB1 |
| PKM | BGN | DCN | RPS16 |
| ANXA2 | MYH9 | COL14A1 | VAT1 |
| ACTN1 | SERPINC1 | CSPG4 | SDCBP |
| F2 | FBLN1 | HSP90B1 | VDAC1 |
| MFGE8 | DCN | IQGAP1 | RAB8A |
| BGN | EDIL3 | ACTN1 | CAP1 |
| FBLN2 | TNC | DARS1 | ITGA5 |
| VCL | GPI | VCL | DSTN |
| PROS1 | LMNA | CAT | PFN1 |
| F10 | LOC506828 | MYH10 | CRP |
| F5 | LOC528040 | COPB2 | C8A |
| ANXA5 | COL6A1 | PRDX1 | MYL12B |
| COL15A1 | EZR | QSOX1 | RPS9 |
| LF | COL14A1 | HSPA8 | TKT |
| COL6A2 | LOC506828 | COL15A1 | TKT |
| TUBB5 | POSTN | HMCN2 | COPA |
| HSP90AA1<br>HSP90AB1<br>APOE<br>LOC506828<br>COL6A1<br>PGAM1 | ALDOA<br>COL6A2<br>ANXA6<br>TUBB5<br>ANXA4<br>PRDX1 | COL14A1<br>EEF2<br>AP2B1<br>GARS1<br>VCP<br>VPS35 | FNDC1<br>CAPNS1<br>NID2<br>LOC100336868<br>MYOF<br>GANAB |

