## Supplemental Figures for "Dynamic Mechanical Loading Reprograms Meniscus Cell-Derived Extracellular Vesicles to Enhance Their Regenerative Potency"

**Supplementary Information**

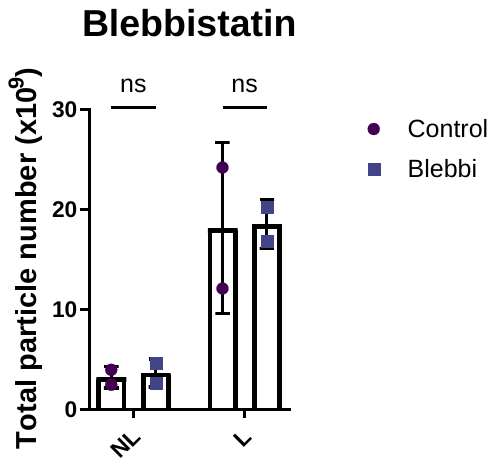

**Figure S1:** Total EV yield from non-loading (NL) and dynamic loading (L) MFCs treated with blebbistatin(−), an inhibitor of actomyosin contractility (n=2 biological replicates).

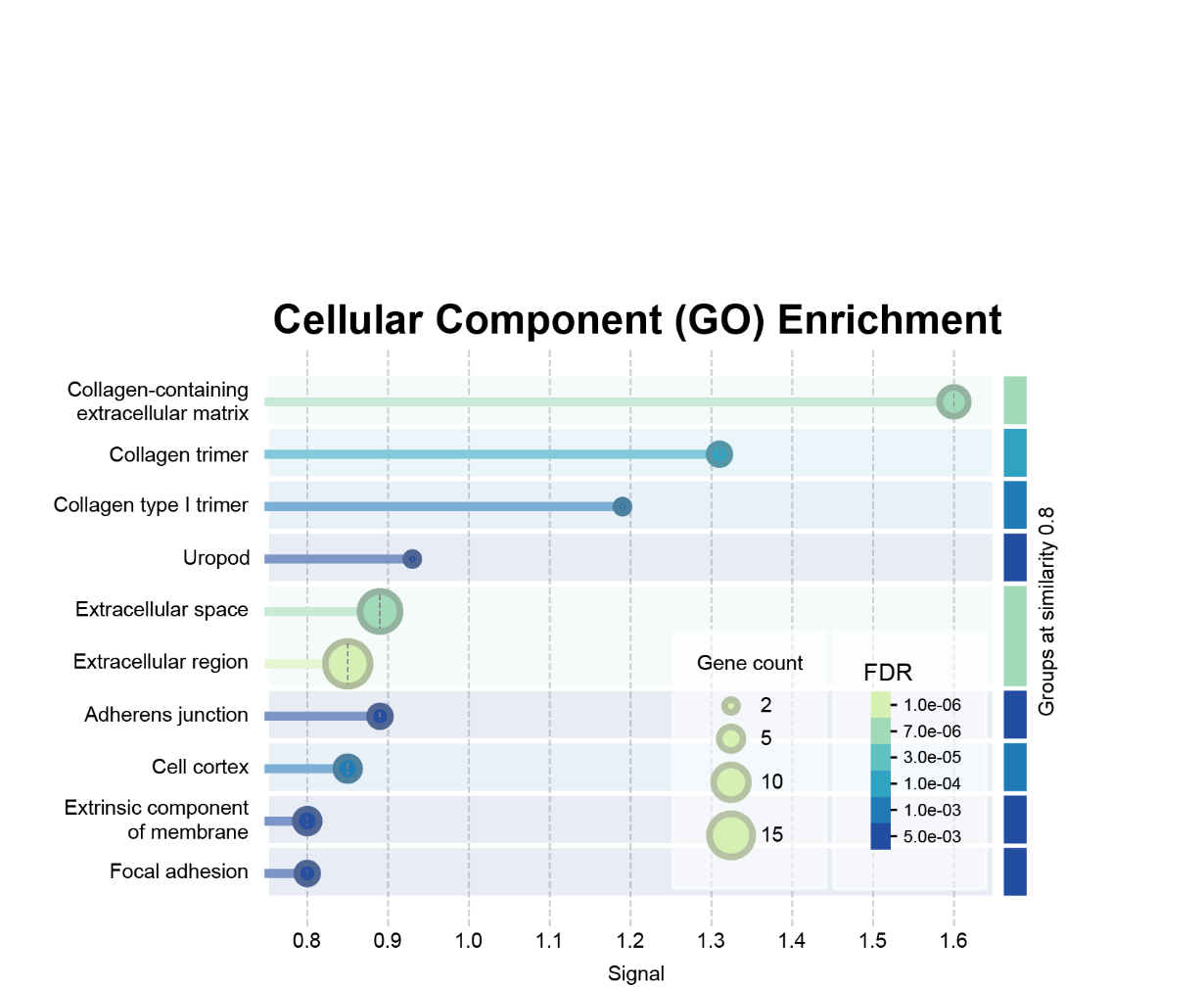

**Figure S2:** Gene Ontology (GO) enrichment analysis of the top 30 upregulated proteins in dynamic mechanical loading (3dL) EVs compared to non-loading (NL) EVs.

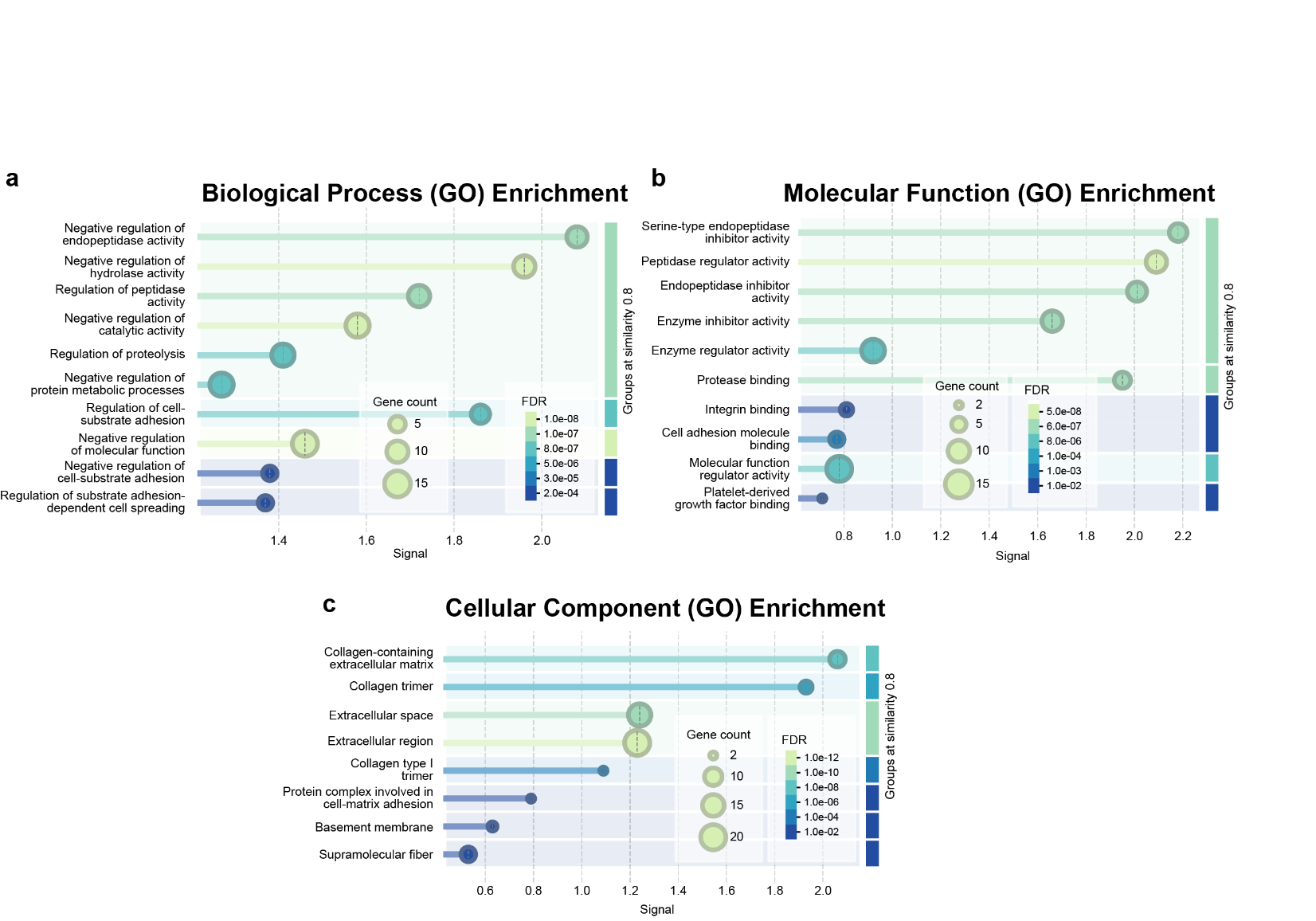

**Figure S3:** Gene Ontology (GO) enrichment analyses of the top 30 proteins identified in non-loading (NL) EVs. Analyses include (a) biological processes, (b) molecular functions, and (c) cellular components.

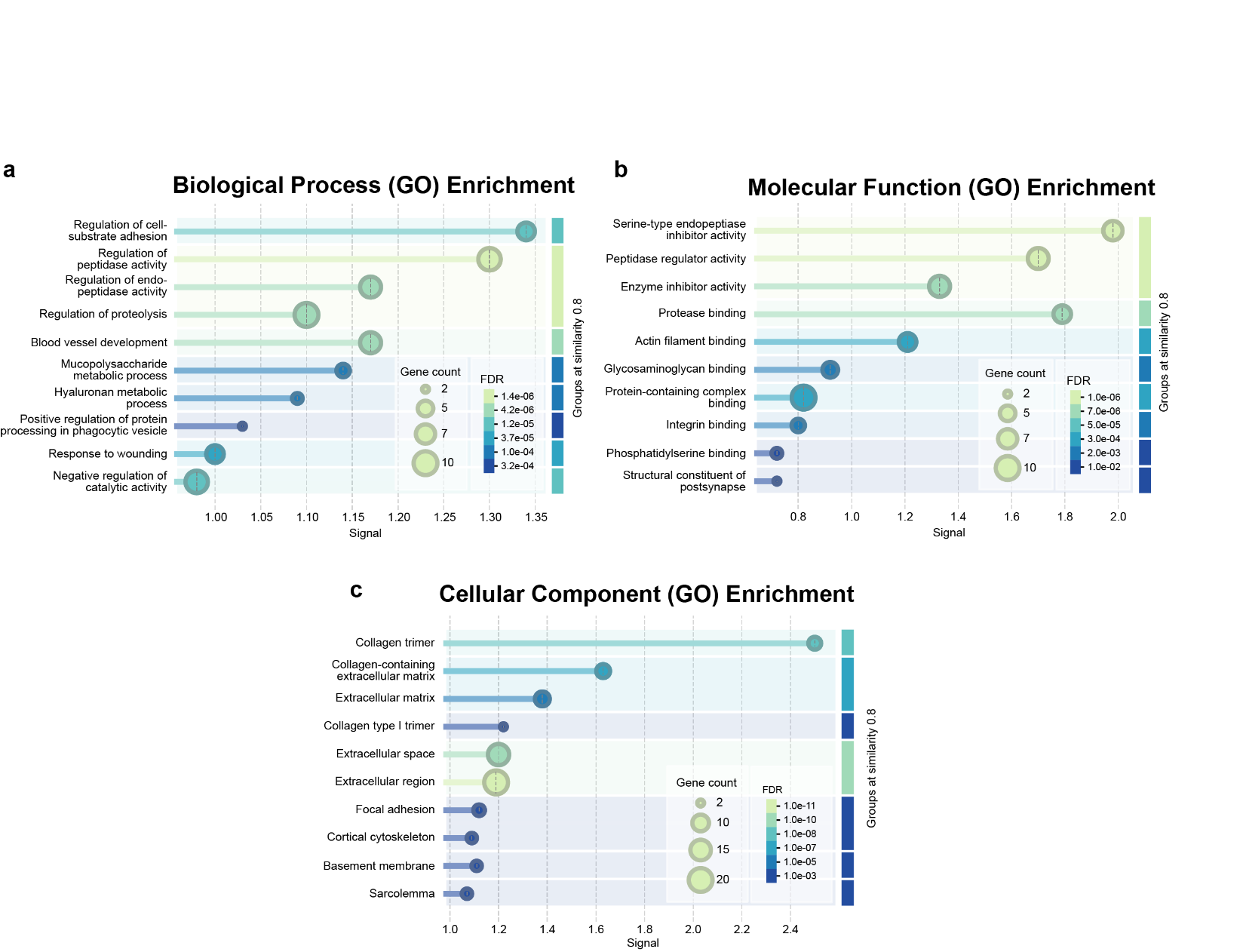

**Figure S4:** Gene Ontology (GO) enrichment analyses of the top 30 proteins identified in mechanically loaded (3dL) EVs. Analyses include (a) biological processes, (b) molecular functions, and (c) cellular components.

**Supplementary Table 1:** Representative top 50 proteins identified by liquid chromatography–tandem mass spectrometry (LC–MS/MS).

| **Top 50 proteins in**  **NL-EVs** | **Top 50 Proteins in**  **3dL-EVs** | **Top 50 proteins upregulated in 3dL-EVs vs. NL-EVs** | **Top 50 proteins only found in 3dL-EVs** |
| --- | --- | --- | --- |
| PLG  A2M  TNC  ADIPOQ  ITIH2  LOC506828  ACTN4  HBB  COL6A3  COL12A1  VTN  COL1A1  POSTN  ACTB  ITIH1  AFP  INHCA  ITIH3  SERPINC1  LOC528040  FBLN1  APOA1  FLNA  COL1A2  MYH9  LOC506828  VIM  TLN1  PKM  ANXA2  ACTN1  F2  MFGE8  BGN  FBLN2  VCL  PROS1  F10  F5  ANXA5  COL15A1  LF  COL6A2  TUBB5  HSP90AA1  HSP90AB1  APOE  LOC506828  COL6A1  PGAM1 | TNC  PLG  MFGE8  FN1  COL1A1  ACTB  ACTN4  ANXA2  ITIH2  A2M  VIM  ITIH1  COL12A1  COL6A3  ADIPOQ  ANXA5  COL1A2  LOC506828  INHCA  ITIH3  ACTN1  VCL  GSN  HBB  SERPINH1  FLNA  PKM  COL15A1  BGN  MYH9  SERPINC1  FBLN1  DCN  EDIL3  TNC  GPI  LMNA  LOC506828  LOC528040  COL6A1  EZR  COL14A1  LOC506828  POSTN  ALDOA  COL6A2  ANXA6  TUBB5  ANXA4  PRDX1 | CDH1  MFGE8  ANXA6  LMNA  ANXA5  ANXA4  ANXA2  GPI  MSN  SERPINH1  PDIA3  GAPDH  PCOLCE  VIM  CA2  TNC  EZR  THBS2  ATP5F1B  EEF1A1  DYNC1H1  NELL2  TNC  COL1A2  COL1A1  HRG  ACTB  ANXA1  DCN  COL14A1  CSPG4  HSP90B1  IQGAP1  ACTN1  DARS1  VCL  CAT  MYH10  COPB2  PRDX1  QSOX1  HSPA8  COL15A1  HMCN2  COL14A1  EEF2  AP2B1  GARS1  VCP  VPS35 | FN1  GSN  EDIL3  ALDOA  ITIH4  MMP2  NID1  SERPINE1  VCAN  HTRA1  MST1  NT5E  ALDOC  MYO1C  ATP1A1  ANXA11  PGK1  ACAN  CLEC3B  PNP  FSCN1  ITGAV  GNAI2  RTN4  YWHAH  LOC784932  SLC1A5  ITGB1  RPS16  VAT1  SDCBP  VDAC1  RAB8A  CAP1  ITGA5  DSTN  PFN1  CRP  C8A  MYL12B  RPS9  TKT  TKT  COPA  FNDC1  CAPNS1  NID2  LOC100336868  MYOF  GANAB |
